# A pumpless and tubeless microfluidic device enables extended *in vitro* development of *Cryptosporidium parvum*

**DOI:** 10.1101/2024.07.07.602413

**Authors:** Samantha Gunasekera, Benjamin Thierry, Edward Cheah, Brendon King, Paul Monis, Jillian M. Carr, Abha Chopra, Mark Watson, Mark O’Dea, Una Ryan

**Affiliations:** Harry Butler Institute, College of Environmental and Life Sciences, Murdoch University, Murdoch 6150, Western Australia, Australia; Future Industries Institute, University of South Australia, Adelaide 5095, South Australia, Australia; South Australian Water Corporation, Adelaide 5000, South Australia, Australia; College of Medicine and Public Health, Flinders University, Flinders Health and Medical Research Institute, Bedford Park 5042, South Australia, Australia; Immunology and Infectious Diseases, Murdoch University, Murdoch 6150, Western Australia, Australia

**Keywords:** *Cryptosporidium*, gut-on-chip, HCT-8 cells, fluid shear stress, *in vitro*

## Abstract

The enteric parasite *Cryptosporidium* remains a treatment challenge for drinking water utilities globally due to its resistance to chlorine disinfection. However, the lack of an *in vitro* culture system for *Cryptosporidium* that is both cost-effective and reliable remains a key bottleneck in *Cryptosporidium* research. Here we report that the microfluidic culture of HCT-8 cells under fluid shear stress enables the extended development of *Cryptosporidium parvum*. Specifically, the growth of *C. parvum* in a user-friendly pumpless microfluidic device was assessed using immunofluorescence assays, scanning electron microscopy and quantitative PCR, which revealed that development peaked at six days post-infection but continued for ten days in total. Oocysts produced within the microfluidic device were infective to fresh HCT-8 monolayers, however these oocysts were only present at low levels. We anticipate that such microfluidic approaches will facilitate a wide range of *in vitro* studies on *Cryptosporidium* and may have the potential to be further developed as a routine infectivity assessment tool for the water industry.

## 1. Introduction

*Cryptosporidium* is an enteric waterborne pathogen and the leading cause of moderate-to-severe diarrhoea in young children in Sub-Saharan Africa and South Asia (1, 2). *Cryptosporidium* has been attributed to ∼12.9 million disability-adjusted life-years (DALYs) lost globally in children under five years of age (3). There is currently no vaccine and very limited treatment options available for cryptosporidiosis (4). Despite the significant global disease burden of cryptosporidiosis, affordable and robust *in vitro* culture systems for *Cryptosporidium* that support complete development remain inaccessible to most laboratories, which is recognised as a critical bottleneck in the development of better therapeutic interventions (5, 6).

Due to the small size of *Cryptosporidium* oocysts (∼5 µm), high shedding rates, low infectious dose, environmental persistence, and resistance to chlorination (7), *Cryptosporidium* was responsible for 76.5% of waterborne outbreaks caused by protozoan parasites globally between 2017 and 2020 (8). Of the > 45 *Cryptosporidium* species described, *C. hominis* and *C. parvum* are the two main species infecting humans and responsible for almost all waterborne outbreaks attributed to this genus (8–11). The accurate assessment of the risk to consumers from *Cryptosporidium* oocysts in drinking water, and the selection of appropriate levels of water treatment, requires quantification of the infectious fraction of oocysts. Relying solely on total oocyst counts or nucleic acid-based detection and identification methods may significantly over-estimate the risk (12). The importance of accurate risk assessment of drinking water sources is increasing as many countries are moving to adopt a tolerable risk of 10^−6^ DALYs per person per year as the health target for drinking water quality (13, 14). Cell culture methods for *Cryptosporidium* infectivity determination using the HCT-8 cell line are the most reliable and widely used (15, 16), and have confirmed that the proportion of infectious oocysts in source water catchments and wastewater treatment plants for drinking water supply can be highly variable (17, 18).

While comparative studies using a wide range of cell lines have indicated that the HCT-8 (human ileocecal colorectal adenocarcinoma) cell line supports superior *Cryptosporidium* development (19), oocyst formation occurs at negligible levels using this system and development will typically cease after 72 hours post-infection (20). Pioneering work utilising stem cell-derived culturing systems have consistently demonstrated that *Cryptosporidium* fertilisation and subsequent oocyst formation can occur under these conditions (21–24), but these approaches remain inaccessible to many labs due to high reagent costs and reliance on primary cells. Bioengineered intestinal models utilising cell lines have also demonstrated complete *C. parvum* development (25, 26). While these studies have been influential for subsequent *in vitro* culture systems for *Cryptosporidium* reported in the literature, they are unsuitable for routine *Cryptosporidium* culture due to the high upfront costs for implementation, complexity of the culturing systems, and inability to scale these systems for diverse applications.

The present study proposes the use of a user-friendly gut-on-chip to extend the current applications of the HCT-8 cell line for the study of *Cryptosporidium* biology, while maintaining ease of use and low costs. The control of fluid flow, and consequent application of fluid shear stress to transformed cells cultured within microfluidic devices, has been frequently documented to change the phenotype of these cells to reflect more physiologically relevant structure and function (27–30). To foster implementation in settings with limited microfluidic experience and facilities, a previously reported pumpless and tubeless microfluidic device was used. In this device, a hydrophilic thread is used to drive the fluid flow, enabling the application of a constant fluid shear stress to cell monolayers as previously validated using the Caco-2 (human colorectal adenocarcinoma) cell line (31). The data presented herein demonstrates that *Cryptosporidium* culture in HCT-8 cells using this technology is cost-effective and reliable, with the potential to be widely used for *Cryptosporidium* infectivity determination.

## 2. Methods

### 2.1 Fabricating microfluidic devices

This study utilised a microfluidic device with a pumpless and tubeless design using a protocol described previously (31). Each device consisted of a glass coverslip bonded to a PDMS layer containing three microchannels (L: 36 mm, W: 1 mm, H: 155 µm). For devices undergoing preparation for scanning electron microscopy, a thicker microscope glass slide was bonded to the PDMS layer instead of cover glass. The total volume of each microchannel was 5.25 µL, and the reservoir volume was 200 µL. Prior to seeding, each microchannel was flushed with 70% (v/v) ethanol, then phosphate buffered saline (PBS) pH 7.4 (Gibco) three times each. A 1% (v/v) Matrigel solution (Corning) diluted with RPMI-1640 (Sigma) was used to coat each microchannel by dispensing the solution through the outlet at 4°C, then incubating for 60 min at 37°C and 5% (v/v) CO_2_ in a humidified incubator.

### 2.2 HCT-8 cell culture inside the microfluidic device

The HCT-8 cell line (human ileocecal colorectal adenocarcinoma, ATCC CCL-244) was routinely cultured in RPMI-1640 supplemented with 10% (v/v) FCS (Bovogen), 1% (v/v) penicillin-streptomycin (Sigma), 2 mM L-glutamine (Sigma), 15 mM HEPES (Sigma), pH 7.2. The cell line was confirmed negative for Mycoplasma prior to experiments using a protocol described previously (32). Cells were harvested from culture flasks using 0.25% (w/v) trypsin-EDTA (Sigma) and seeded into Matrigel-coated microchannels via the outlet at a concentration of 2 × 10^5^ cells per microchannel. The microfluidic devices were incubated overnight at 37°C with 5% (v/v) CO_2_ without the thread to facilitate cell adherence. The reservoir of each microchannel was then filled with RPMI-1640 supplemented as described above. To initiate fluid flow, a pre-sterilised hand-cut 10 mm spunlace thread (70% rayon, 30% polyester, Ebos Healthcare Australia) was inserted into the outlet of each microchannel using sterile, fine-tipped forceps. Cell monolayers were then grown to 90% confluence under shear stress (0.02 dyn cm^−2^) for downstream experiments. For transwell comparisons, the HCT-8 cell line was seeded at a concentration of 5 × 10^5^ cells per well into 24-well plates and grown to 90% confluence.

### 2.3 Excystation pre-treatment of oocysts, sporozoite purification and infection of HCT-8 cells

The *C. parvum* (Iowa-IIaA17G2R1) oocysts were obtained from BioPoint Pty Ltd (Sydney, Australia) and the subtype was confirmed using a nested PCR targeting the 60-kilodalton glycoprotein (*gp60*) locus described previously (33, 34), with minor modifications to reagent and primer concentrations reported elsewhere (35). The *C. parvum* oocysts underwent excystation pre-treatment as described previously (36). Oocysts were resuspended in 20 µL of infection medium per microchannel, which comprised RPMI-1640 supplemented with 2 mM L-glutamine, 5.6 mM glucose, 0.02% (w/v) bovine bile, 15 mM HEPES, 0.6 µM folic acid, 7.3 µM 4-aminobenzoic acid, 2.1 µM calcium pantothenate, 50 µM L-ascorbic acid, 2.5 µg mL^−1^ amphotericin B, 1% (v/v) penicillin-streptomycin (all from Sigma) and 1% (v/v) FCS (Bovogen), pH 7.2. For experiments designed to detect new oocyst production within the microfluidic device, purified sporozoites were used for infections. The excystation pre-treated oocysts were resuspended in 1 mL infection medium and then incubated at 37°C for 30 min, followed by filtration with a 2 µm syringe filter (Whatman). The syringe filter was rinsed with an additional 2 mL infection medium, then all filtrate containing purified sporozoites was centrifuged at room temperature at 3,200 x *g* for 20 min and resuspended in a pre-calculated volume of infection medium.

For infection, the spunlace thread was removed from each outlet, then excystation pre-treated oocysts were applied to HCT-8 monolayers via the outlet at concentrations of 1 × 10^4^ oocysts or 4 × 10^5^ sporozoites per microchannel in a volume of 20 µL of infection medium. The devices were then incubated at 37°C and 5% (v/v) CO_2_ for four hours without the thread, after which fluid flow was re-initiated by inserting a fresh thread into the outlet of each microchannel. Infection medium from each microchannel and reservoir was replaced at 24 h intervals. For transwell comparison experiments, excystation pre-treated *C. parvum* oocysts were applied to HCT-8 monolayers and cultured in 24-well plates at a concentration of 1 × 10^4^ oocysts per well.

### 2.4 Parasite enumeration using immunofluorescence assays (Sporo-Glo Ab) and quantitative PCR

For experiments designed to quantify parasite multiplication over time, a total of 16 microchannels were seeded with HCT-8 cells, and 15 were infected with excystation pre-treated *C. parvum* oocysts as described above. The uninfected microchannel seeded with HCT-8 cells functioned as a negative control for immunofluorescence assays and as a no template control (NTC) for quantitative PCR assays. Three microchannels were prepared for immunofluorescence assays at 48 h intervals throughout the experiment. At each time-point, the thread was removed from each outlet, submerged in 20 µL of infection medium and stored at 4°C. The infection medium inside each microchannel and reservoir was then collected via the outlet and stored in 1.5 mL tubes at 4°C. For transwell comparisons, three wells per plate were allocated to each time-point. The microfluidic devices and transwell plates were prepared for immunostaining as described previously (37). A fluorescein-labelled anti-*Cryptosporidium* polyclonal antibody (1X Sporo-Glo, Waterborne Inc) was administered to each microchannel directly from the outlet, or to each transwell, at a volume of 100 µL, and then incubated in the dark at room temperature for 60 min. After immunostaining, the remaining Sporo-Glo was aspirated, then the cell monolayers were rinsed four times using PBS and imaged using a Nikon Eclipse TS100 inverted fluorescence microscope.

Following image acquisition, cell monolayers were harvested from each microchannel and each transwell using 0.25% (w/v) trypsin-EDTA (Sigma). The trypsin-EDTA was inactivated with an equal volume of infection medium and transferred to 1.5 mL tubes. All samples collected throughout experiments from harvested monolayers, threads and residual infection medium from each time-point were centrifuged at 20,000 x *g* for 10 min, and the supernatant was removed from each tube, leaving ∼20 µL remaining. All samples were freeze-thawed five times using alternating 60 second incubation periods in liquid nitrogen and 120 second incubation periods on a heating block set to 100°C. On the final thaw, each sample was boiled at 100°C for 5 min and then the lysate was stored at −20°C until ready for use.

A quantitative PCR assay targeting a *Cryptosporidium*-specific gene encoding a C-type lectin protein was used to measure parasite load over time. The primer set (forward primer: EVF1 5′-GAA CTG TAC AGA TGC TTG GGA GAA T and reverse primer: EVR1 5′-CCTT CGT TAG TTG AAT CCT CTT TCC A) amplified a 150 bp product and was used with a minor groove binding (MGB) probe (5′JOE – CTT GGA GCT CGT ATC AG – MGBNFQ) that was specific to *C. parvum* (38). DNA was amplified using a CFX96 Touch Real-Time PCR Detection System (Bio-Rad) in a 10 µL reaction containing 1X GoTaq Probe qPCR Master Mix (Promega), 800 nm EVF1 forward primer, 800 nm EVR1 reverse primer, 250 nm *C. parvum* MGB probe and 2 µL lysate diluted 1:100. The reactions were carried out under the following cycling conditions: 95°C for 2 minutes, then 45 cycles of 95°C for 30 seconds and 60°C for 60 seconds. A standard curve was produced using a serially diluted gBlocks dsDNA Gene Fragment (Integrated DNA Technologies) that was synthesised to be homologous to the region of interest, with the AT-rich regions modified for increased GC content. Quantitative PCR data was acquired and analysed using the CFX Manager software (Bio-Rad) and converted to oocyst equivalents on the basis that the gene is single copy (39, 40) and there are four haploid sporozoites per oocyst.

### 2.5 Preparation of microfluidic devices for scanning electron microscopy

A combination of uninfected HCT-8 monolayers and *C. parvum*-infected monolayers were prepared for imaging using scanning electron microscopy. The microchannels were coated with 1% (v/v) Matrigel, seeded with HCT-8 cells and grown to confluence as described above. Each microchannel was then infected with 1.0 × 10^5^ excystation pre-treated *C. parvum* oocysts. The microchannels were rinsed three times with PBS and fixed overnight in 2.5% (v/v) glutaraldehyde diluted in 100 mM HEPES buffer (both from Sigma), pH 7.4. For *Cryptosporidium*-infected microchannels, fixation occurred at 48 h post-infection, whereas uninfected HCT-8 monolayers underwent fixation once the monolayer reached 90% confluence. The glutaraldehyde was rinsed off using 100 mM HEPES buffer, keeping the entire device fully submerged in aqueous solution. The PDMS layer of the microfluidic device was removed from the microscope slide using a scalpel blade, then each sample was subject to graded ethanol and hexamethyldisilazane dehydration. Samples were then sputter coated with 5 nm platinum. Morphological data were acquired using a Zeiss Supra 55VP field emission scanning electron microscope using the in-lens secondary electron detector and an accelerating voltage of 5 kV. The *C. parvum* life cycle stages were identified using morphological criteria described previously (41).

### 2.6 Detection of new oocysts produced *in vitro*

For experiments designed to detect new oocysts produced *in vitro*, six microchannels were seeded with HCT-8 cells as described above. Three microchannels were infected with purified *C. parvum* sporozoites and the remaining microchannels were designated as the negative control for immunofluorescence assays. All infection medium inside each microchannel and reservoir was collected in 1.5 mL tubes at 24 h intervals for seven days in total. All samples were centrifuged at 8,000 x *g* for 3 min and washed twice with PBS. Samples were then prepared for immunostaining using EasyStain (BioPoint Pty Ltd) using the supplier’s instructions and imaged using a Nikon Eclipse 80i Fluorescence Microscope. To determine whether the *in vitro* produced oocysts were infectious, the infection medium from one microchannel at three days post-infection underwent excystation pre-treatment before application to a fresh HCT-8 monolayer. After 48 hours, the HCT-8 monolayer was prepared for immunostaining using Sporo-Glo as described above and imaged using a Nikon Eclipse TS100 inverted fluorescence microscope.

## 3. Results

This study reports the *in vitro* development of *C. parvum* in HCT-8 cell monolayers grown within the microfluidic device for up to 10 days. HCT-8 cells cultured at 0.02 dyn cm^−2^ exhibited well-developed microvilli-like structures (Figure 1 B-C). Sampling was undertaken at 48 h intervals from the cell culture medium within the microchannel, the absorbent thread initiating fluid flow, and the cell monolayer harvested from each microchannel. Quantitative PCR values for these samples were pooled together as the total parasite load (referred to as oocyst equivalents) at each time-point (Figure 2A).

**Figure 1.**
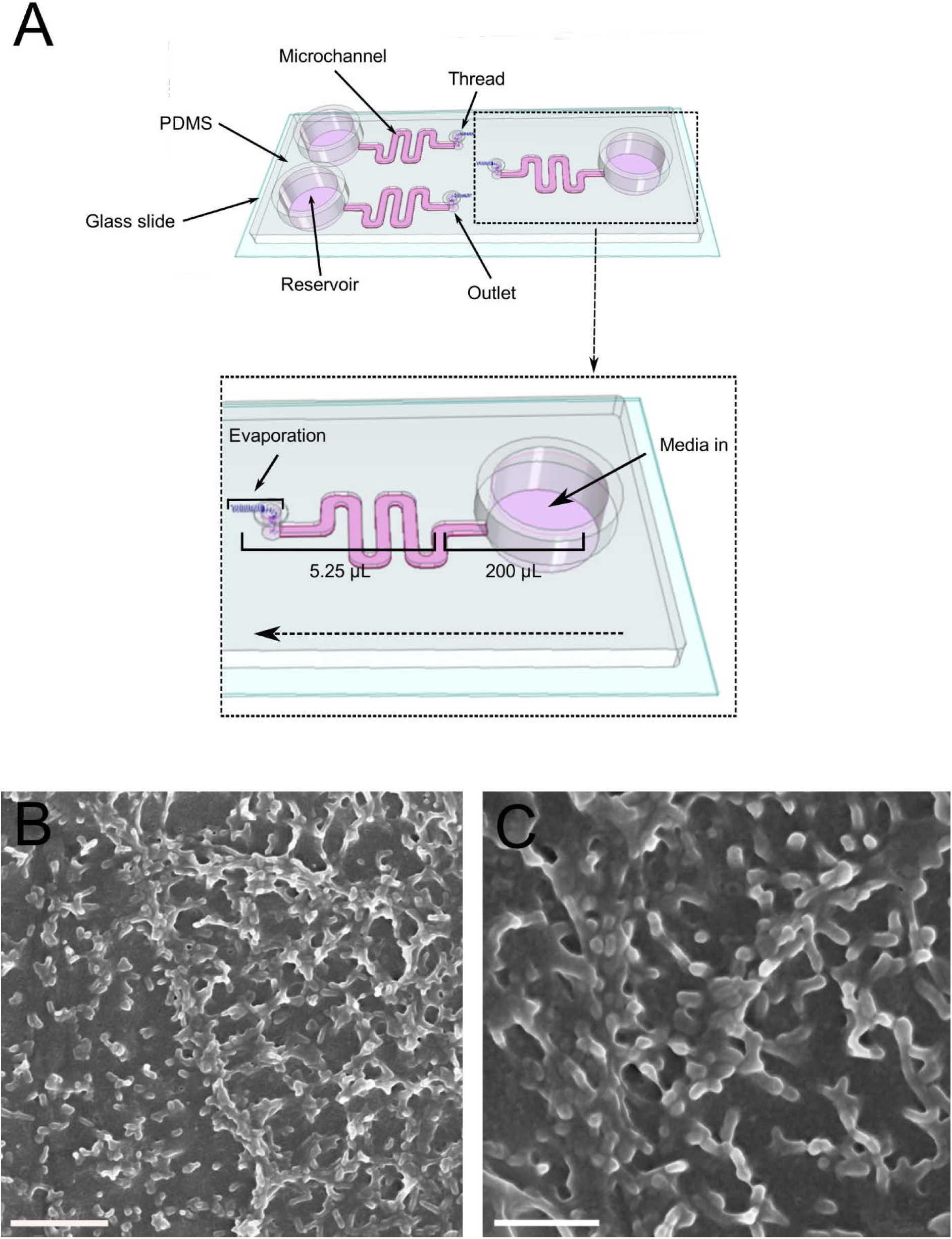
HCT-8 cells cultured under fluid shear stress within the microfluidic device enables formation of microvilli-like structures. A: Schematic image of the microfluidic device indicating the location of each microchannel, reservoir, outlet and thread placement (diagram not drawn to scale). B-C: Scanning electron micrographs of the HCT-8 cell monolayer. HCT-8 cells were grown in Matrigel-coated microchannels under shear stress at 0.02 dyn cm^−2^ until 90% confluent (up to five days). The cell monolayers were then fixed with glutaraldehyde, subjected to graded ethanol and chemical dehydration, and coated with 5 nm platinum. Data were acquired using a field emission scanning electron microscope. Representative images are shown. Scale bars: B = 2 µm, C = 1 µm.

**Figure 2.**
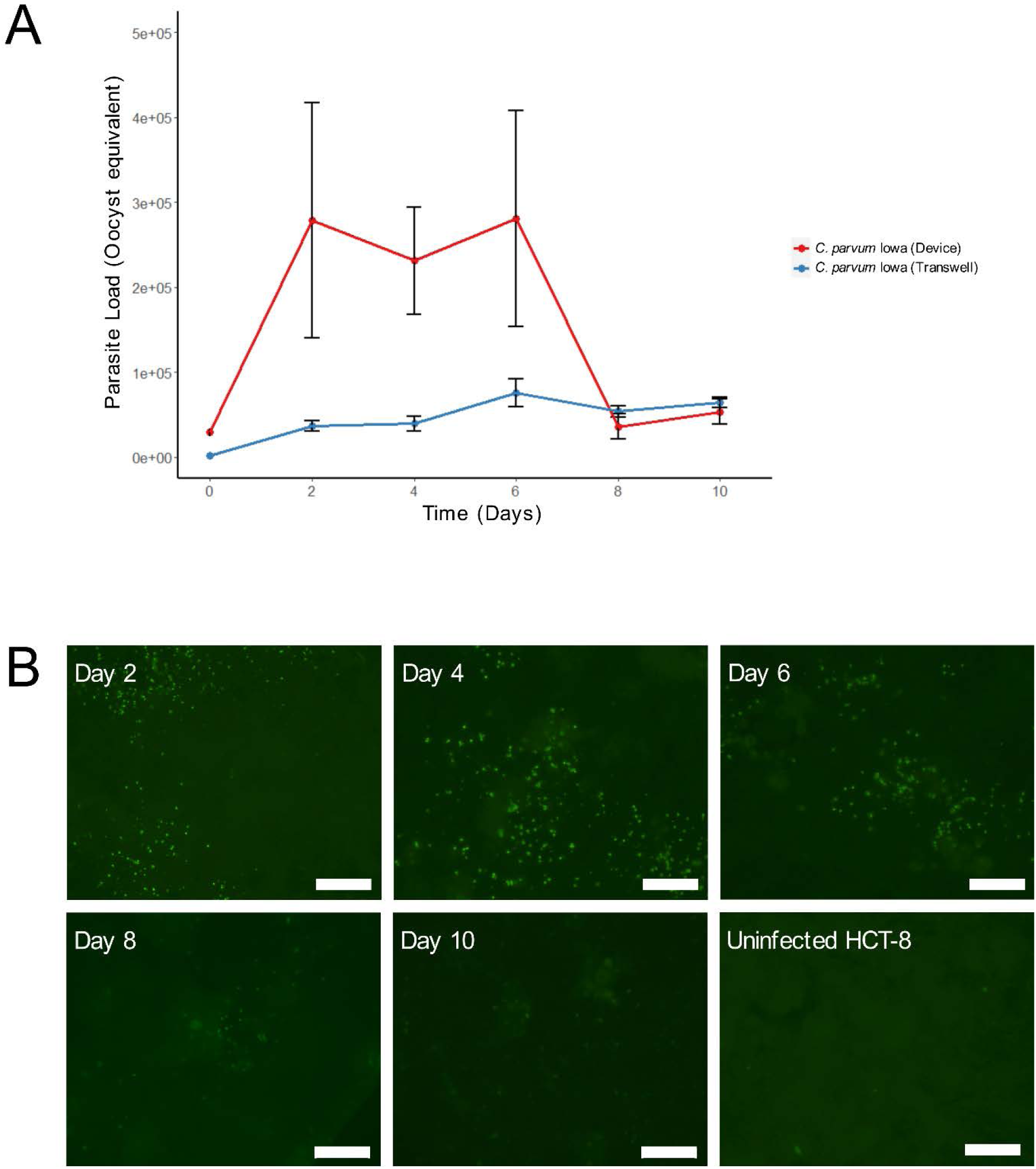
The fluid shear stress conditions of the microfluidic device enable long term *C. parvum* development. A: Quantitative PCR of *C. parvum* infection of the HCT-8 monolayer cultured within the microfluidic device vs. transwells. HCT-8 cells were infected with excystation pre-treated *C. parvum* oocysts and the infection medium, thread and monolayer were sampled at 48 h intervals. The overall parasite load was calculated from a standard curve and normalised for the total volume of the original, undiluted lysate. Results represent means ± standard deviations from *n* = 3 independent experiments performed in triplicate under fluid shear stress in the microfluidic device and *n* = 1 independent experiments performed in triplicate under static conditions in transwell plates. B: HCT-8 cells cultured under fluid shear stress infected with excystation pre-treated *C. parvum* Iowa oocysts, then fixed and stained with Sporo-Glo (fluorescein-labelled polyclonal anti-*Cryptosporidium* antibody) at each time-point from two to ten days post-infection (scale bars = 100 µm).

The quantitative PCR data was supported by immunofluorescence assays using the Sporo-Glo polyclonal antibody, which indicated that foci of infection were present in the HCT-8 monolayer at each time-point up to ten days post-infection (Figure 2B). No immunostaining was detected in the uninfected HCT-8 monolayers. The number of foci visible in the monolayers declined rapidly from eight to ten days post-infection, which coincided with what was observed from the quantitative PCR data.

Scanning electron micrographs of the *Cryptosporidium*-infected microfluidic devices at two days post-infection demonstrated the presence of all common life cycle stages, including both asexual and sexual phases of development. The scanning electron micrographs indicated the presence of sporozoites, trophozoites, meronts and merozoites attached to the host cell surface. The presence of microgamonts and macrogamonts were visible at this time-point (Figure 3). The invasive life cycle stages (sporozoites, merozoites) were observed rarely. Trophozoites and meronts were regularly observed. Of the sexual stages, macrogamonts were more frequently observed than microgamonts.

**Figure 3.**
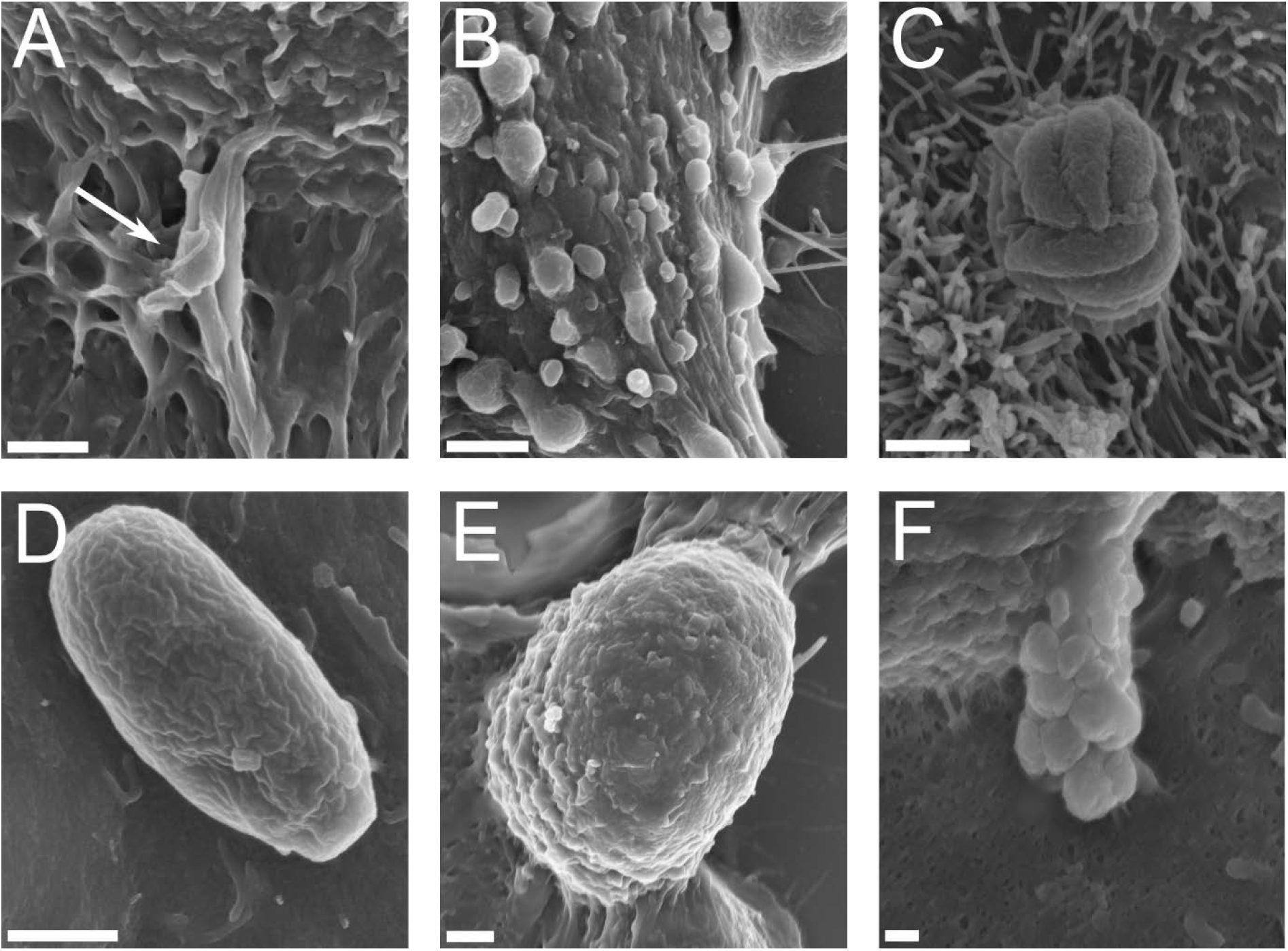
Scanning electron micrographs of *C. parvum* infecting the HCT-8 cell line grown under fluid shear stress conditions at two days post-infection. A: Sporozoite invading the host cell monolayer (indicated by arrowhead), B: Trophozoites adhered to the host cell monolayer appearing as spherical structures on the cell surface, C: Meront adhered to the host cell, D: Merozoite attached lengthwise to the apical surface of the host cell, E: Macrogamont attached to the host cell, F: Microgamont hanging from a stalk with multiple microgametes visible. Scale bars: A-E = 1 µm, F = 200 nm.

A commercially obtained antibody that binds to *Cryptosporidium* oocyst walls (EasyStain) was used with 4’,6-diamidino-2-phenylindole (DAPI) to identify the presence of newly formed *Cryptosporidium* oocysts within the microfluidic device. The EasyStain immunofluorescence assay indicated that oocysts were present in low numbers in the cell culture medium of the *C. parvum* infected microchannels from three to ten days post-infection (Figure 4), and oocyst shells were present in the cell culture medium at one day post-infection. The oocysts that were detected in the cell culture medium often did not stain with DAPI (Figure 4). Oocyst shells were observed, but rarely and in very low numbers when sampled immediately post-filtration and at two days post-infection. When the cell culture medium from a *C. parvum*-infected microchannel at three days post-infection was applied to a fresh HCT-8 monolayer, foci of infection were observed at 48 h post-infection of the passaged culture, indicating that the oocysts produced *in vitro* within the microfluidic HCT-8 cell cultures were infectious to a fresh HCT-8 cell monolayer (Figure 4).

**Figure 4.**
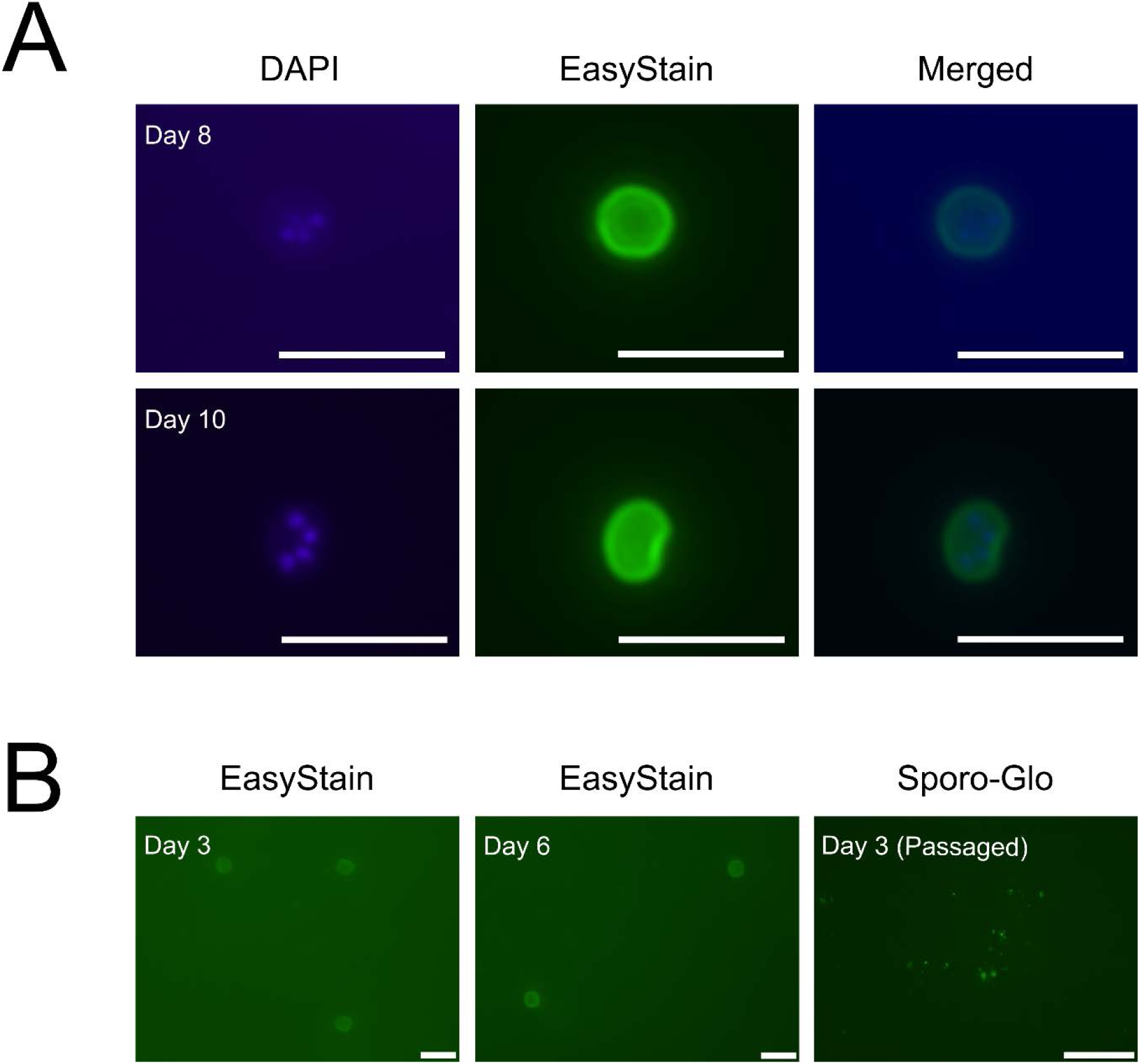
The microfluidic device enables formation of *C. parvum* oocysts that are infectious to fresh HCT-8 monolayers. A: Contents of one microfluidic device from eight days and ten days post-infection stained with DAPI, EasyStain, and merged image for both time-points. B: Left two images are the contents of one microfluidic device from three days and six days post-infection and stained with EasyStain for both time-points. The image on the right is an HCT-8 monolayer with the contents of one microchannel at three days post-infection, applied to the fresh monolayer and cultured in transwells for 48 h before immunostaining with Sporo-Glo. Scale bars: A = 10 µm, B = 10 µm (left), 100 µm (right).

## 4. Discussion

This study demonstrated the *in vitro* development of *C. parvum* using the HCT-8 cell line for up to ten days using a pumpless and tubeless microfluidic device. Our data indicated an increase in parasite load over time, with the infection peaking at six days post-infection and then declining rapidly from day eight onwards. The quantitative PCR data was supported by immunofluorescence data indicating robust infection up to six days post-infection, followed by low numbers of foci of infection at eight days post-infection which declined further by ten days post-infection. Scanning electron micrographs demonstrated a large amount of heterogeneity of both asexual and sexual developmental stages after two days of infection under fluid shear stress conditions. Under static conditions, *in vitro* development of *C. parvum* has been previously observed to be largely asexual at the same time-point (35, 41).

The seminal work of English et al. (2022) provided insights into the infection kinetics of *C. parvum*, where it was demonstrated that the parasite must undergo three cycles of asexual replication before sexual development will occur (42). Within the transwell plate, fertilisation is very limited which results in cessation of *C. parvum* infection after three days post-infection (20, 43). The findings of the present work suggest that one cycle of successful sexual replication occurred within the microfluidic device, and excystation of oocysts produced *in vitro* occurred within the microchannel. The data suggested subsequent cycles of asexual development occurring after three days post-infection. The loss of reliability of the culturing system after six days of infection may coincide with a possible second round of sexual development resulting in a diminished number of oocysts produced *in vitro* to sustain infection within the microfluidic device, and may also reflect the ageing of the HCT-8 host cells. While the present study provided evidence of the production of new and viable oocysts *in vitro* at three days post-infection, we present insufficient evidence to indicate that the fertilisation block observed in conventional HCT-8 cell culture is completely circumvented with the microfluidic gut-on-chip, given that the new oocysts were present in relatively low numbers. The presence of oocysts that would not stain with DAPI but were still infective to new cultures was unsurprising due to an unclear relationship between oocyst viability and permeability to vital dyes (44, 45).

In the *in vivo* environment, gut epithelial cells are exposed to a wide variety of mechanical stimuli that result in the application of shear stress, including peristaltic muscle contraction, intraluminal fluid flow, and rhythmic extension and contraction of the intestinal villi (46). The application of fluid shear stress on gut epithelial cell lines cultured *in vitro* within microfluidic devices has been demonstrated to alter their phenotype, translating to a more physiologically relevant representation of the *in vivo* structure and function. Such phenotypical changes elicited by fluid shear stress in gut epithelial cells include increased cell height, improved density of F-actin networks, enhanced microvilli and tight junction formation, and elevated mucin production (27–29, 47–49).

The majority of gut-on-chip microfluidic devices require integration of an external peristaltic pump and tubing to control perfusion of cell culture medium through the microchannels (50, 51). In practice, these are prone to leakages (52), and therefore suboptimal for culturing *Cryptosporidium* due to the elevated biological hazard this creates. The pumpless and tubeless microfluidic device utilised in the present study minimises this risk and can be easily integrated into existing workflows that do not routinely use microfluidics due to its simplified design. These devices can be easily prepared at a low upfront cost within basic microfabrication facilities (< $1 AUD per microfluidic device). Given that many labs that study the *in vitro* development of *Cryptosporidium* already have robust HCT-8 transwell-based culturing systems, this microfluidic device can integrate seamlessly into existing workflows and extend the boundaries of what can currently be achieved with the transwell system.

Currently, the costs associated with routine *Cryptosporidium* infectivity testing limit their widespread usage within the water industry, at an estimated cost per sample of ∼$400 (AUD) for the *Cryptosporidium* culturing component alone. Forgoing *Cryptosporidium* infectivity testing due to cost limitations can lead to failures in accurately quantifying the risk of *Cryptosporidium* oocyst detections in source water and challenges in meeting health-based targets for water quality. Using microfluidic gut-on-chips instead of transwells for *Cryptosporidium* infectivity testing could contribute to minimising the reagent costs due to the small volume of the microchannel (5.25 µL) and reservoir (200 µL), and results in a total cost reduction of ∼17%. For laboratories wanting to apply this technology for routine *Cryptosporidium* infectivity studies, it could provide a more affordable system to determine the proportion of infectious oocysts recovered from drinking water and wastewater supplies using either the focus detection method with Sporo-Glo (37) or by combining with quantitative PCR (38). In the future, the microfluidic device described in the present study may form part of an integrated assay to quantitate *Cryptosporidium* oocyst viability and identify species and subtype to better inform water quality risk assessments.

## 5. Conclusions

We have shown that an easy-to-use, inexpensive microfluidic device supports the growth of all commonly observed life cycle stages of *C. parvum* within HCT-8 cells for up to ten days. We anticipate that this approach will make studies on *Cryptosporidium* infectivity, life cycle, host-parasite interactions and high-throughput drug assays more accessible to both researchers and industry and contribute to the advancement of our understanding of the biology of *Cryptosporidium*.

## 6. Declarations

### 6.1 Patient consent statement

No human participants, human data or human tissue requiring ethics approval was used in this study.

### 6.2 Consent for publication

Not applicable.

### 6.3 Availability of data and materials

Further information and requests for materials should be directed to the corresponding author.

### 6.6 Authors’ contributions

SG: Conceptualisation, methodology, formal analysis, investigation, writing – original draft, writing – review and editing, visualisation. BT: Conceptualisation, methodology, resources, writing – review & editing, funding acquisition. EC: Methodology, resources, visualisation. BK: Conceptualisation, methodology, writing – review and editing, funding acquisition. PM: Conceptualisation, methodology, writing – review and editing, funding acquisition. JMC: Conceptualisation, writing – review and editing, funding acquisition. AC: Methodology, resources, supervision. MW: Methodology, resources, supervision, writing – review and editing. MOD: Conceptualisation, supervision, writing – review and editing, funding acquisition. UR: Conceptualisation, methodology, resources, writing – original draft, writing – review and editing, supervision, project administration, funding acquisition.

## 6.7 Acknowledgements

We thank Dr Andrew Ball from Water New South Wales, Dr Nicholas Crosbie from Melbourne Water, Dr Paul Fisher from Seqwater and Dr Leon van der Linden from South Australian Water Corporation for their role as industry partners and contribution to funding. We thank Ms Frances Brigg for performing Sanger sequencing and A/Prof Peta Clode for technical advice in performing the sample preparation for scanning electron microscopy. The authors acknowledge use of the facilities of Microscopy Australia at the Centre for Microscopy, Characterisation & Analysis, The University of Western Australia, a facility funded by the University, State and Commonwealth Governments.

